# Lipid Biosynthesis Perturbation Impairs Endoplasmic Reticulum-Associated Degradation

**DOI:** 10.1101/2022.12.09.519544

**Authors:** Samantha M. Turk, Christopher J. Indovina, Danielle L. Overton, Avery M. Runnebohm, Cade J. Orchard, Ellen M. Doss, Kyle A. Richards, Courtney Broshar Irelan, Mahmoud M. Daraghmi, Connor G. Bailey, Jacob M. Miller, Julia M. Niekamp, Samantha K. Gosser, Mary E. Tragesser-Tiña, Kieran P. Claypool, Sarah M. Engle, Bryce W. Buchanan, Kelsey A. Woodruff, James B. Olesen, Philip J. Smaldino, Eric M. Rubenstein

## Abstract

The relationship between lipid homeostasis and protein homeostasis (proteostasis) is complex and remains incompletely understood. We conducted a screen for genes required for efficient degradation of *Deg1*-Sec62, a model aberrant translocon-associated substrate of the endoplasmic reticulum (ER) ubiquitin ligase Hrd1, in *Saccharomyces cerevisiae*. This screen revealed that *INO4* is required for efficient *Deg1*-Sec62 degradation. *INO4* encodes one subunit of the Ino2/Ino4 heterodimeric transcription factor, which regulates expression of genes required for lipid biosynthesis. *Deg1*-Sec62 degradation was also impaired by mutation of genes encoding several enzymes mediating phospholipid and sterol biosynthesis. The degradation defect in *ino4*Δ yeast was rescued by supplementation with metabolites whose synthesis and uptake are mediated by Ino2/Ino4 targets. Stabilization of a panel of substrates of the Hrd1 and Doa10 ER ubiquitin ligases by *INO4* deletion indicates ER protein quality control is generally sensitive to perturbed lipid homeostasis. Further, loss of *INO4* sensitized yeast to proteotoxic stress, suggesting a broad requirement for lipid homeostasis in maintaining proteostasis. Abundance of the ER ubiquitin-conjugating enzyme Ubc7 was reduced in the absence of *INO4*, consistent with a model whereby perturbed lipid biosynthesis alters the abundance of critical protein quality control mediators, with broad consequences for ER proteostasis. A better understanding of the dynamic relationship between lipid homeostasis and proteostasis may lead to improved understanding and treatment of several human diseases associated with altered lipid biosynthesis.

## INTRODUCTION

Proteome maintenance is crucial for eukaryotic life. Dedicated mechanisms to destroy aberrant or overabundant proteins are present in many cellular compartments. A substantial proportion of protein turnover at the endoplasmic reticulum (ER) is accomplished through ER-Associated Degradation (ERAD; reviewed in (Mehrtash and Hochstrasser 2019; Christianson and Carvalho 2022)). In ERAD, ubiquitin ligases transfer ubiquitin from ubiquitin-conjugating enzymes to aberrant or overabundant proteins, which are subsequently degraded by the 26S proteasome. The two major ERAD ubiquitin ligases in *Saccharomyces cerevisiae* are the highly conserved multipass transmembrane enzymes, Hrd1 and Doa10 (Hampton *et al*. 1996; Plemper *et al*. 1999; Swanson *et al*. 2001). Hrd1 functions with the soluble ubiquitin-conjugating enzyme Ubc7 and, to a lesser extent, Ubc1 and Ubc6 (Plemper *et al*. 1999; Bays *et al*. 2001; Lips *et al*. 2020). Doa10 functions with two ubiquitin-conjugating enzymes, Ubc7 and the transmembrane protein Ubc6 (Swanson *et al*. 2001). Ubc7 is tethered to the ER membrane and stabilized by the transmembrane protein Cue1 (Biederer *et al*. 1997; Bazirgan and Hampton 2008; Metzger *et al*. 2013).

Hrd1 and Doa10 differentially target ERAD substrates based on the location and nature of the substrates’ degrons, or degradation signals. Hrd1 ubiquitylates soluble and integral membrane proteins with ER luminal degrons and proteins that clog translocons, which transfer proteins into or across the ER membrane (Carvalho *et al*. 2006; Gauss *et al*. 2006; Rubenstein *et al*. 2012; Runnebohm *et al*. 2020b). Conversely, Doa10 recognizes soluble and integral membrane proteins with cytosolic degrons (Huyer *et al*. 2004; Ravid *et al*. 2006; Metzger *et al*. 2008). Both enzymes target proteins with intramembrane degrons (Sato *et al*. 2009; Habeck *et al*. 2015). While Hrd1 resides exclusively in the ER membrane, Doa10 is also found in the contiguous inner nuclear membrane (INM), where it ubiquitylates proteins with nucleoplasmic degrons (Deng and Hochstrasser 2006). Additional ubiquitin ligases contribute to the degradation of proteins at the ER and INM. The ubiquitin ligases Ubr1 and Ltn1 contribute to ERAD (Stolz *et al*. 2013; Crowder *et al*. 2015; Arakawa *et al*. 2016), and the INM Asi complex and anaphase-promoting complex mediate turnover of aberrant or overabundant INM proteins (Foresti *et al*. 2014; Khmelinskii *et al*. 2014; Koch *et al*. 2019). Finally, the metalloprotease Ste24 contributes to degradation of translocon-clogging proteins via a mechanism that is partially redundant with Hrd1-mediated ubiquitylation (Ast *et al*. 2016; Runnebohm *et al*. 2020b).

While much has been uncovered about the molecular mechanisms of ERAD, a comprehensive characterization of genetic requirements for ER protein degradation remains incomplete (Runnebohm *et al*. 2020b). We conducted a genome-wide, growth-based reporter screen to identify potential yeast genes required for turnover of a model translocon-clogging substrate of Hrd1. This screen revealed that *INO4*, which encodes one subunit of a heterodimeric transcription factor that regulates several genes encoding lipid-biosynthetic enzymes, is required for efficient translocon quality control (TQC) substrate degradation. We found TQC is broadly sensitive to perturbations in phospholipid and sterol biosynthesis. Further, a panel of model Hrd1 and Doa10 substrates bearing luminal, intramembrane, and cytosolic degrons were stabilized by *INO4* deletion, and yeast with defects in phospholipid or sterol synthesis were sensitive to conditions associated with aberrant protein production. The abundance of Ubc7, which is broadly required for ERAD, was reduced in *ino4*Δ yeast, suggesting a possible mechanism for disrupted ERAD. Taken together, our results indicate that altered lipid homeostasis broadly and profoundly impairs ER proteostasis. Several metabolic, muscular, cardiac, and neurodegenerative diseases are associated with perturbed lipid synthesis (Li *et al*. 2006; Mitsuhashi and Nishino 2011; Arendt *et al*. 2013; Kosicek and Hecimovic 2013; Kim *et al*. 2016; Bargui *et al*. 2021). Altered lipid homeostasis may impair ER protein degradation in individuals with these disorders.

## METHODS

### Yeast and Plasmid Methods

Yeast were cultured at 30°C in standard growth medium (Guthrie and Fink 2004). Plasmids were introduced into yeast via lithium acetate transformation (Guthrie and Fink 2004). See **Table S1** for yeast strains used in this study. See **Table S2** for plasmids used in this study.

To generate pVJ490 (a plasmid containing *P*_*GAL4*_*-DEG1-SEC62-HIS3* and *natMX4* as independent genes that could be amplified as a single PCR product for genomic integration), a 1314-bp EagI fragment containing *natMX4* from pAG25 (alias pVJ132) (Goldstein and Mccusker 1999) was inserted into the EagI site of pVJ477 (Watts *et al*. 2015), which possessed *P*_*GAL4*_*-DEG1-SEC62-HIS3*. Orientation of the *natMX4* fragment was confirmed by NcoI digestion.

To generate the query strain VJY355 for Synthetic Genetic Array analysis (Tong *et al*. 2001), a cassette containing *P*_*GAL4*_*-DEG1-SEC62-HIS3* and *natMX4* flanked by 50 bp of DNA homologous to sequence upstream and downstream of the *DOA10* open reading frame was PCR-amplified from pVJ490 using primers VJR264 and VJR265 (see **Table S3** for primers used in this study). This PCR product was introduced to Y7049 (haploid MATα query strain; alias VJY338 (Tong and Boone 2007)), and integration was confirmed by three-primer PCR at the 5’ and 3’ ends of the *doa10*Δ::*P*_*GAL4*_*-DEG1-SEC62-HIS3*:*natMX4* locus using primers VJR46, VJR82, and VJR260 (5’ end) and VJR11, VJR107, and VJR249 (3’ end). To confirm integration of the cassette at a single locus, VJY355 was mated with nourseothricin-sensitive MAT<u>a</u> haploid yeast. Sporulation of the mated diploid was induced, and 2:2 segregation of nourseothricin resistance:sensitivity was observed, consistent with integration of the *P*_*GAL4*_*-DEG1-SEC62-HIS3*:*natMX4* cassette at a single locus.

To generate VJY951 (*hrd1*Δ:: *kanMX4 ino4*Δ::*kanMX4*), MATα *hrd1*Δ::*kanMX4* yeast (VJY478) were mated with MAT<u>a</u> *ino4*Δ::*kanMX4* yeast (VJY474). Mated *HRD1*/*hrd1*Δ::*kanMX4 INO4*/*ino4*Δ::*kanMX4* heterozygous diploids were induced to undergo sporulation, and spores were separated by microdissection. Candidate double mutant yeast were selected on the basis of 2:2 segregation of G418 resistance:sensitivity, and *HRD1* and *INO4* genotypes were verified by PCR using primers VJR70, VJR163, and VJR259 (to distinguish *HRD1* and *hrd1*Δ::*kanMX4*) and primers VJR371, VJR372, and VJR259 (to distinguish *INO4* and *ino4*Δ::*kanMX4*).

For supplementation experiments, yeast were cultured (from inoculation until cell harvest and cycloheximide chase) in media containing 500 μM inositol, 2 mM ethanolamine, and 2 mM choline. For the inositol limitation experiment, cells were cultured to mid-exponential growth in medium containing inositol, washed six times in inositol-free medium (prepared using yeast nitrogen base without amino acids and inositol), and incubated in inositol-free medium for 5 h. Serial dilution growth assays were performed as described (Watts *et al*. 2015).

### Screen of Yeast Deletion and Hypomorphic Allele Collections

Screening of the yeast genome was performed as described in (Tong *et al*. 2001). VJY355 (a MATα *his3*Δ*1* query strain possessing *P*_*GAL4*_*-DEG1-SEC62-HIS3* and *natMX4* at the *DOA10* locus) was mated with the haploid yeast MAT<u>a</u> *his3*Δ*1* knockout and DAmP (Decreased Abundance by mRNA Perturbation) libraries of non-essential and essential genes, respectively (Winzeler *et al*. 1999; Giaever *et al*. 2002; Breslow *et al*. 2008). Each 96-well plate of the yeast knockout and DAmP collections includes a blank well (no yeast); *hrd1*Δ yeast (VJY22; positive control) were spiked into the blank well of each plate as a positive control for *Deg1*-Sec62-His3 stabilization. Following serial transfer using 96-prong pinners and culture of yeast on a series of selective media, a library of haploid MAT<u>a</u> yeast expressing *Deg1*-Sec62-His3 and possessing knockout or hypomorphic alleles of each gene represented in the knockout and DAmP collections was generated. These yeast were transferred to 96-well plates possessing synthetic complete media and cultured for 48 h at 30°C. 40 μl of each culture were transferred to 96-well plates containing 160 μl of selective media lacking histidine. The OD_595_ for each strain was recorded at the beginning and end of an 11-h incubation period at 30°C using an iMark Microplate Absorbance Reader (Bio-Rad). A detailed outline of the SGA procedure is included in **Table S4**.

### Cycloheximide Chase

Cycloheximide chase experiments were performed as previously described (Buchanan *et al*. 2016). Briefly, mid-exponential phase yeast cultured at 30°C were concentrated to 2.5 OD_600_ units/ml in fresh synthetic defined medium and maintained at 30°C. Cycloheximide was added to each culture (final concentration 250 μg/ml). 2.4-OD_600_ aliquots were harvested immediately after cycloheximide addition and at indicated time points and were added to stop mix containing sodium azide (final concentration 10 mM) and bovine serum albumin (final concentration 0.25 mg/ml). Samples were maintained on ice until the end of the chase, at which point all yeast were lysed.

### Cell Lysis

Unless otherwise indicated, yeast were lysed using the alkaline lysis method as previously described (Kushnirov 2000; Watts *et al*. 2015). 2.4 – 2.5 OD_600_ units were harvested and suspended in 200 μl of 0.1 M NaOH, followed by incubation at room temperature for 5 min and pelleting by centrifugation. Pellets were resuspended in 1X Laemmli sample buffer and boiled at 100°C for 5 min. Insoluble material was pelleted by high-speed centrifugation, and the soluble fraction (supernatant) was retained for electrophoresis.

For the experiment presented in **Figure 4B**, yeast were lysed using a trichloracetic acid (TCA) lysis procedure as previously described (Loayza and Michaelis 1998). 2.4 OD_600_ units of yeast were harvested and suspended in 0.26 M NaOH and 0.13 M Δ-mercaptoethanol, followed by incubation on ice for 15 min. TCA (final concentration 5%) was added to cell suspensions to precipitate proteins, followed by centrifugation at 4°C. Pellets were resuspended in TCA sample buffer (3.5% SDS, 0.5 M DTT, 80 mM Tris, 8 mM EDTA, 15% glycerol, 0.1 mg/ml bromophenol blue) and heated to 37°C for 30 min. Insoluble material was pelleted by centrifugation (18,000 x *g* for 1 min), and the soluble fraction (supernatant) was retained for analysis by SDS-PAGE.

### Western Blotting

Following separation by SDS-PAGE, proteins were transferred to polyvinylidene difluoride (PVDF) membrane via wet transfer at 20 V for 1 h at 4°C. Membranes were blocked in 5% skim milk suspended in Tris-buffered saline (TBS; 50 mM Tris, 150 mM NaCl) at room temperature for 1 h or overnight at 4°C. Antibody incubations were performed in 1% skim milk suspended in TBS with 1% Tween 20 (TBS/T) for 1 h at room temperature followed by three five-min washes in TBS/T. The following primary antibody dilutions were used: mouse anti-HA.11 (Clone 16B12; BioLegend) at 1:1,000; mouse anti-GFP (Clone JL-8; Clontech) at 1:1,000; and mouse anti-Pgk1 (Clone 22C5D8; LifeTechnologies) at 1:20,000 – 1:40,000. Mouse primary antibodies were followed by incubation with AlexaFlour-680-conjugated rabbit anti-mouse secondary antibody (LifeTechnologies) at 1:20,000 – 1:40,000. Rabbit primary antibodies were followed by incubation with DyLight-800-conjugated goat anti-rabbit secondary antibody (Invitrogen). *Deg1*\*-Sec62 and *Deg1*-Vma12 possess two copies of *Staphylococcus aureus* Protein A epitope, which interacts non-specifically with mammalian immunoglobulins (Hjelm *et al*. 1972) and were directly detected using the AlexaFlour-680-conjugated rabbit anti-mouse antibody. PVDF membranes were imaged with the Odyssey CLx IR Imaging System (Li-Cor) and analyzed with ImageStudio software (Li-Cor).

## RESULTS

### Genome-wide screen to identify genes required for degradation of model Translocon Quality Control substrate

We conducted a reporter-based screen to identify genes required for efficient degradation of the model TQC substrate *Deg1*-Sec62. Fusing His3 to the C-terminus of *Deg1*-Sec62 (**Figure 1A**) allows selection of degradation-defective mutant yeast lacking endogenous *HIS3*. Yeast unable to degrade *Deg1*-Sec62-His3 exhibit histidine prototrophy ((Watts *et al*. 2015), **Figure 1B**).

**Figure 1.**
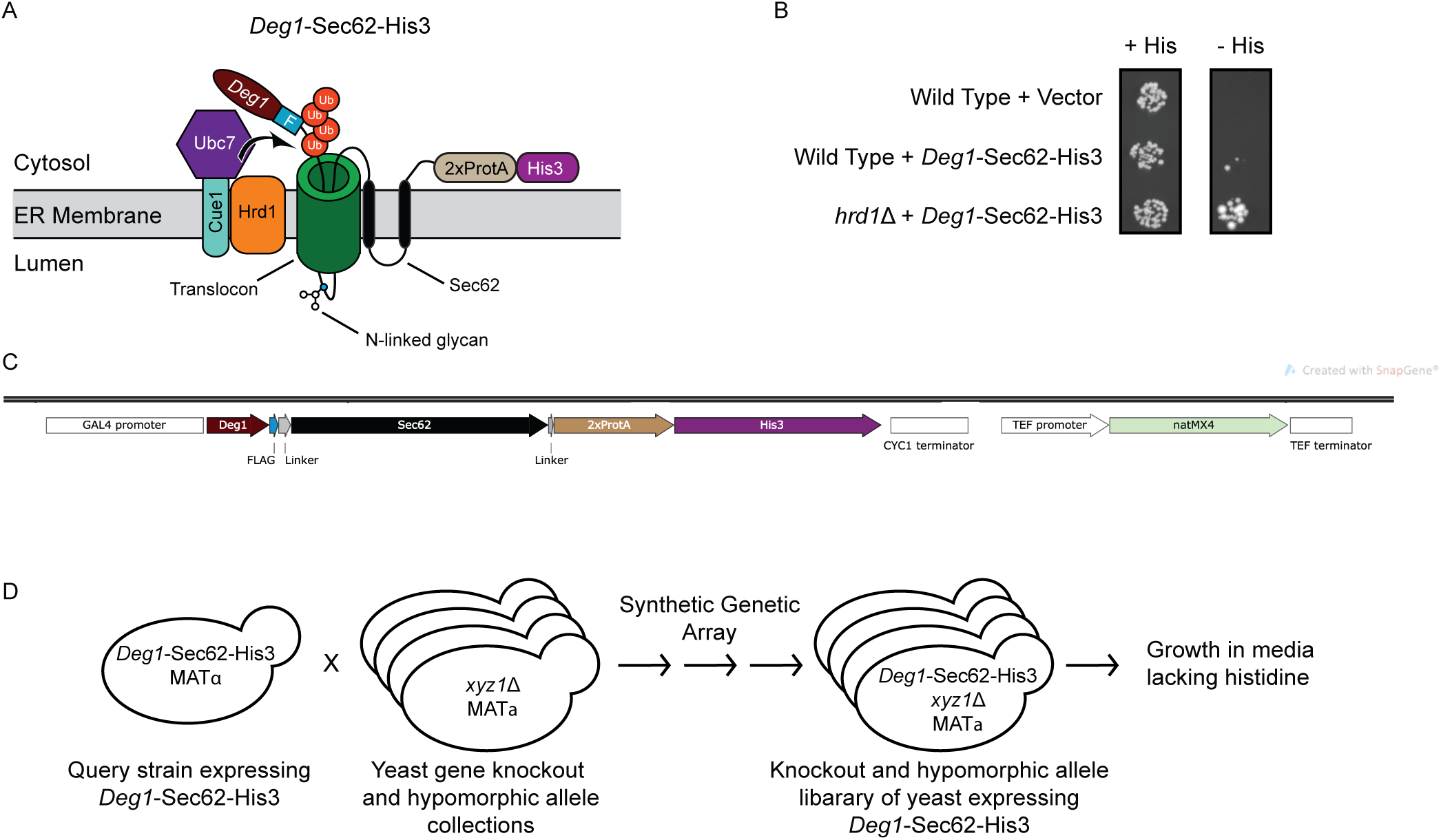
Screen for genes required for degradation of a model translocon-clogging protein. **(A)** Schematic of *Deg1*-Sec62-His3 following aberrant translocon engagement. Following integration of the two transmembrane segments of Sec62, the N-terminal tail of the fusion protein loops into and persistently engages (i.e. clogs) the translocon (Rubenstein *et al*. 2012). Upon clogging, *Deg1*-Sec62(±His3) is ubiquitylated by Hrd1 and Ubc7 (which is anchored at the ER membrane by Cue1). *Deg1*-Sec62-His3 possesses, in sequence, *Deg1* (the N-terminal 67 amino acids from the yeast transcriptional repressor MATα2), a FLAG epitope (F), Sec62, two copies of *Staphylococcus aureus* Protein A (2xProtA), and the His3 enzyme. Ub, ubiquitin. **(B)** Yeast of the indicated genotypes transformed with an empty vector or a plasmid encoding *Deg1*-Sec62-His3 were spotted onto media containing or lacking histidine (His). **(C)** *DOA10* locus of the query strain used for the genome-wide screen. *DOA10* was replaced with a cassette containing *Deg1-Sec62-His3* and *natMX4* as two independent genes, each with its own promoter and transcriptional terminator. **(D)** Overview of genome-wide screen. See text and **Table S4** for details.

A query strain encoding *Deg1*-Sec62-His3 driven by the *GAL4* promoter (**Figure 1C**) was crossed with collections of yeast strains possessing deletions of non-essential genes and hypomorphic alleles of essential genes. Using Synthetic Genetic Array (SGA) technology (Tong *et al*. 2001), a library of ∼6,000 unique mutant strains harboring *Deg1*-Sec62-His3 was generated (**Figure 1D**). Under some conditions (e.g. when ER translocation is impaired), *Deg1*-Sec62 is converted from a Hrd1 substrate into a Doa10 substrate (Rubenstein *et al*. 2012). Therefore, to simplify analysis, the gene encoding *Deg1*-Sec62-His3 was introduced at the *DOA10* locus, replacing *DOA10* in the query strain.

Each mutant strain with *Deg1*-Sec62-His3 was inoculated into liquid media (containing histidine) in a 96-well plate and allowed to incubate at 30°C for 48 h. Equal volumes of each culture were transferred to fresh media lacking histidine and incubated for 11 h at 30°C. The optical density at 595 nm (OD_595_) of each strain was recorded at the beginning and end of the 11 h incubation period. A cutoff for ΔOD_595_ values of 0.079 was selected, resulting in 128 genes encoding proteins with potential roles in *Deg1*-Sec62-His3 degradation (**File S1**). Deletion of *GAL80* (Johnston et al. 1987), which encodes a repressor of the *GAL4* promoter used to drive expression of *Deg1*-Sec62-His3, yielded the highest ΔOD_595_ value. *HRD1, HRD3* (which encodes a Hrd1 cofactor (Hampton *et al*. 1996; Rubenstein *et al*. 2012)), and *UMP1* (which encodes a proteasome assembly factor (Ramos *et al*. 1998)) were identified in this screen, engendering confidence in the power of this analysis to yield bona fide genetic requirements for protein degradation.

Gene Ontology (GO) analysis (www.yeastgenome.org) of the 128 genes revealed significant enrichment of genes linked to processes related to sulfur metabolism (*sulfate assimilation, sulfur compound biosynthetic process, sulfur amino acid metabolic process, toxin biosynthetic process, hydrogen sulfide metabolic process*, and *hydrogen sulfide biosynthetic process*) (**Table S5**). No GO terms relating to function or component were significantly enriched.

A subset of mutants with ΔOD_595_ values at or above the 0.079 cutoff were selected for further evaluation. Naïve yeast with mutations in genes identified in the screen were transformed with plasmids encoding *Deg1*-Sec62-His3 and/or *Deg1*\*-Sec62 for confirmatory reporter-based growth assays and/or biochemical analysis (i.e. cycloheximide chase experiments and western blots), respectively, as indicated in **File S1**. *Deg1*\*-Sec62 possesses mutations that preclude degradation by the Doa10 pathway while still permitting Hrd1-mediated degradation (Johnson *et al*. 1998; Rubenstein *et al*. 2012). Loss of three genes not previously implicated in TQC enhanced stability of *Deg1*\*-Sec62. Deletion of *INO4* and *KAR3* (which encodes a minus-end-directed kinesin) strongly stabilized *Deg1*\*-Sec62, while *SET2* (which encodes a histone methyltransferase) modestly, but reproducibly, delayed *Deg1*\*-Sec62 turnover (**Figure 2A, Figure S1**).

**Figure 2.**
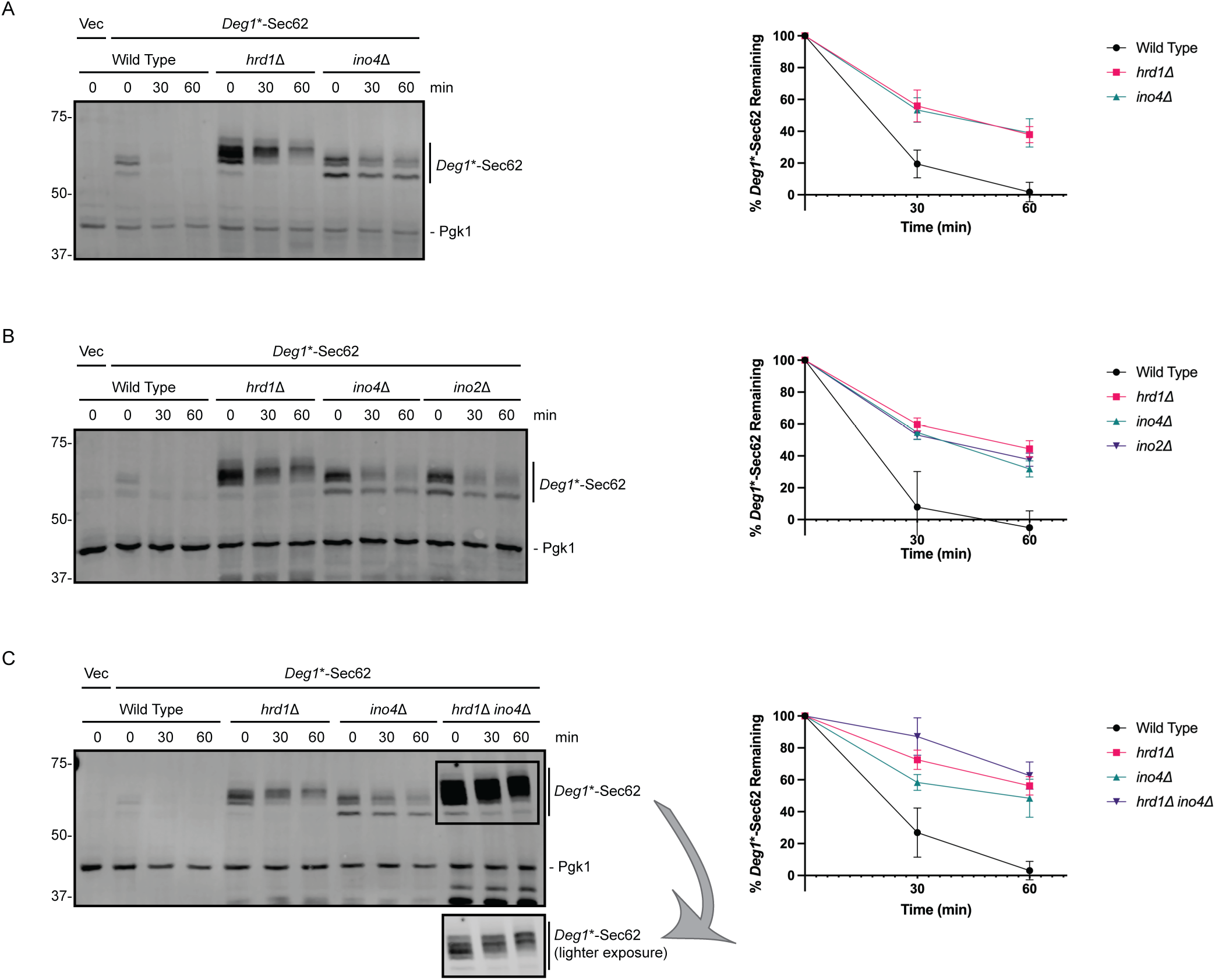
Ino2 and Ino4 are required for *Deg1*\*-Sec62 degradation. Left, Yeast of the indicated genotypes were transformed with a plasmid encoding *Deg1*\*-Sec62 or an empty vector and subjected to cycloheximide chase and western blot analysis to detect *Deg1*\*-Sec62 and Pgk1. Right, quantification of experiments. Error bars represent Standard Error of the Mean of n = 3-5 experiments.

### *INO2* and *INO4* are required for Translocon Quality Control degradation of *Deg1*\*-Sec62

The Ino2/Ino4 heterodimeric transcription factor regulates expression of at least 88 genes, many of which encode enzymes involved in phospholipid synthesis (Ambroziak and Henry 1994; Henry *et al*. 2012). The Ino2/Ino4 complex has not previously been implicated in ERAD. Cycloheximide chase analysis confirmed that loss of *INO4* stabilizes *Deg1*\*-Sec62 to a similar extent as *HRD1* deletion (**Figure 2A**). *Deg1*\*-Sec62 migrates as multiple bands; appearance of higher molecular weight species reflects N-linked glycosylation of the protein, which occurs upon persistent translocon engagement (Rubenstein *et al*. 2012). In addition to slowing degradation of *Deg1*\*-Sec62, *INO4* deletion delays the appearance of higher molecular weight species. Loss of Ino4’s binding partner Ino2 similarly stabilized and delayed modification of *Deg1*\*-Sec62 (**Figure 2B**).

*HRD1* deletion incompletely stabilizes *Deg1*\*-Sec62, consistent with multiple degradation pathways acting on translocon-clogging proteins (Runnebohm *et al*. 2020b). We generated *hrd1*Δ *ino4*Δ yeast to ascertain if *INO4* deletion prevents Hrd1-dependent or -independent degradation. While *Deg1*\*-Sec62 steady state abundance is substantially increased in the double mutant, the rate of *Deg1*\*-Sec62 degradation is only marginally decreased in *hrd1*Δ *ino4*Δ yeast compared to that observed in either single mutant strain (**Figure 2C**). This suggests the impact of *INO2*/*INO4* deletion on *Deg1*\*-Sec62 degradation is largely through modulation of Hrd1-dependent turnover.

### Degradation of *Deg1*\*-Sec62 is sensitive to perturbations in lipid biosynthesis

Supplementation of media with inositol, ethanolamine, and choline (intermediates whose synthesis and uptake are mediated by targets of Ino2/Ino4 (Henry *et al*. 2012)) restored *Deg1*\*-Sec62 degradation (**Figure 3A**). This is consistent with perturbed lipid biosynthesis causing the degradation defect. Ino2/Ino4 regulates expression of genes encoding enzymes mediating multiple branches of phospholipid biosynthesis. We analyzed *Deg1*\*-Sec62 degradation in yeast harboring deletion or hypomorphic alleles of three of these genes: *INO1, CDS1*, and *CHO2*. Ino1 catalyzes the conversion of glucose-6-phosphate to a precursor of inositol and is essential for *de novo* synthesis of phosphatidylinositol derivatives (Donahue and Henry 1981). Cds1 promotes the synthesis of cytidine diphosphate, a precursor of several membrane lipids, including phosphatidylinositol derivatives, cardiolipin, phosphatidylserine, phosphatidylethanolamine, and phosphatidylcholine (Shen *et al*. 1996). Cho2 is required for *de novo* synthesis of phosphatidylcholine (Summers *et al*. 1988; Kodaki and Yamashita 1989). Mutation of *CDS1* or *INO1* strongly stabilized *Deg1*\*-Sec62 (**Figure 3B**). Deletion of *CHO2* also slowed *Deg1*\*-Sec62 turnover (**Figure 3C**).

**Figure 3.**
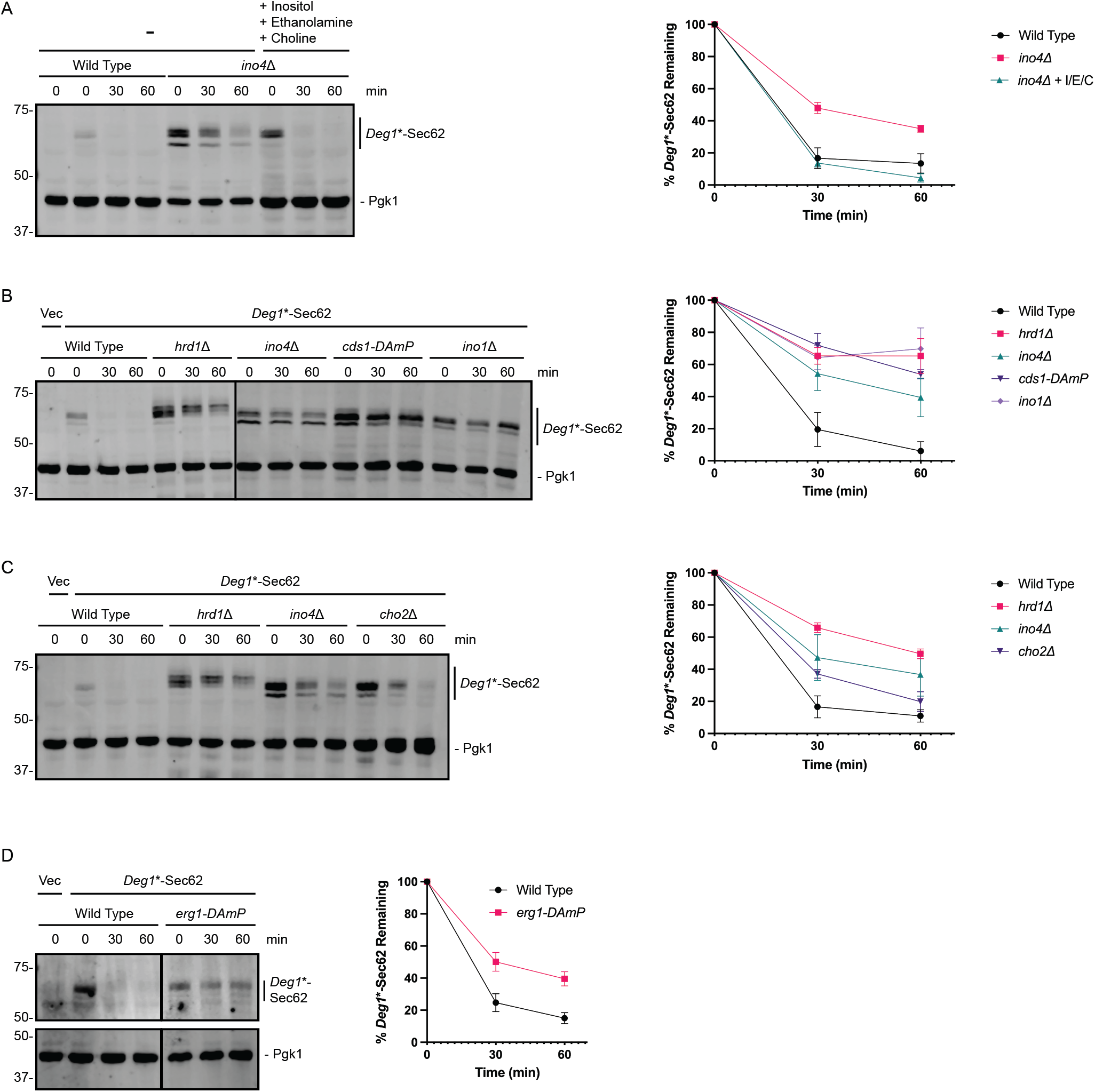
*Deg1*\*-Sec62 degradation is sensitive to perturbations of lipid biosynthesis. Left, Yeast of the indicated genotypes were transformed with a plasmid encoding *Deg1*\*-Sec62 or an empty vector and subjected to cycloheximide chase and western blot analysis to detect *Deg1*\*-Sec62 and Pgk1. Right, quantification of experiments. Error bars represent Standard Error of the Mean of n = 3-5 experiments. For experiment depicted in **(A)**, final three lanes represent yeast supplemented with 500 μM inositol, 2 mM ethanolamine, and 2 mM choline from inoculation through cell harvest and cycloheximide chase.

*ERG1* encodes squalene epoxidase, which mediates an essential step in biosynthesis of ergosterol, the primary sterol in fungal membranes (Jandrositz *et al*. 1991). The ΔOD_595_ value for *erg1-DAmP* yeast in our screen was 0.078, just beyond the 0.079 cutoff. Perturbation of *ERG1*, which is not regulated by Ino2/Ino4, enhanced *Deg1*\*-Sec62 stability (**Figure 3D**). Together, these results indicate *Deg1*\*-Sec62 degradation is broadly sensitive to perturbation in membrane lipid biosynthesis.

### *INO4* deletion causes a generalized ERAD defect and a reduction in Ubc7 abundance

We investigated the requirement of *INO4* for the degradation of a panel of Hrd1 and Doa10 ERAD substrates (**Figure 4A**). Strikingly, *INO4* deletion stabilized an integral membrane Hrd1 substrate bearing an intramembrane degradation signal (HA-Pdr5*, **Figure 4B** (Plemper *et al*. 1998)), a soluble luminal Hrd1 substrate (CPY*-HA, **Figure 4C** (Hiller *et al*. 1996)), an integral membrane Doa10 substrate bearing a cytosolic degron (*Deg1*-Vma12, **Figure 4D** (Ravid *et al*. 2006)), and a soluble, cytosolic Doa10 substrate (*Deg1*-GFP, **Figure 4E** (Lenk and Sommer 2000)). Thus, deletion of *INO4* broadly impairs protein degradation mediated by the ERAD ubiquitin ligases Hrd1 and Doa10.

**Figure 4.**
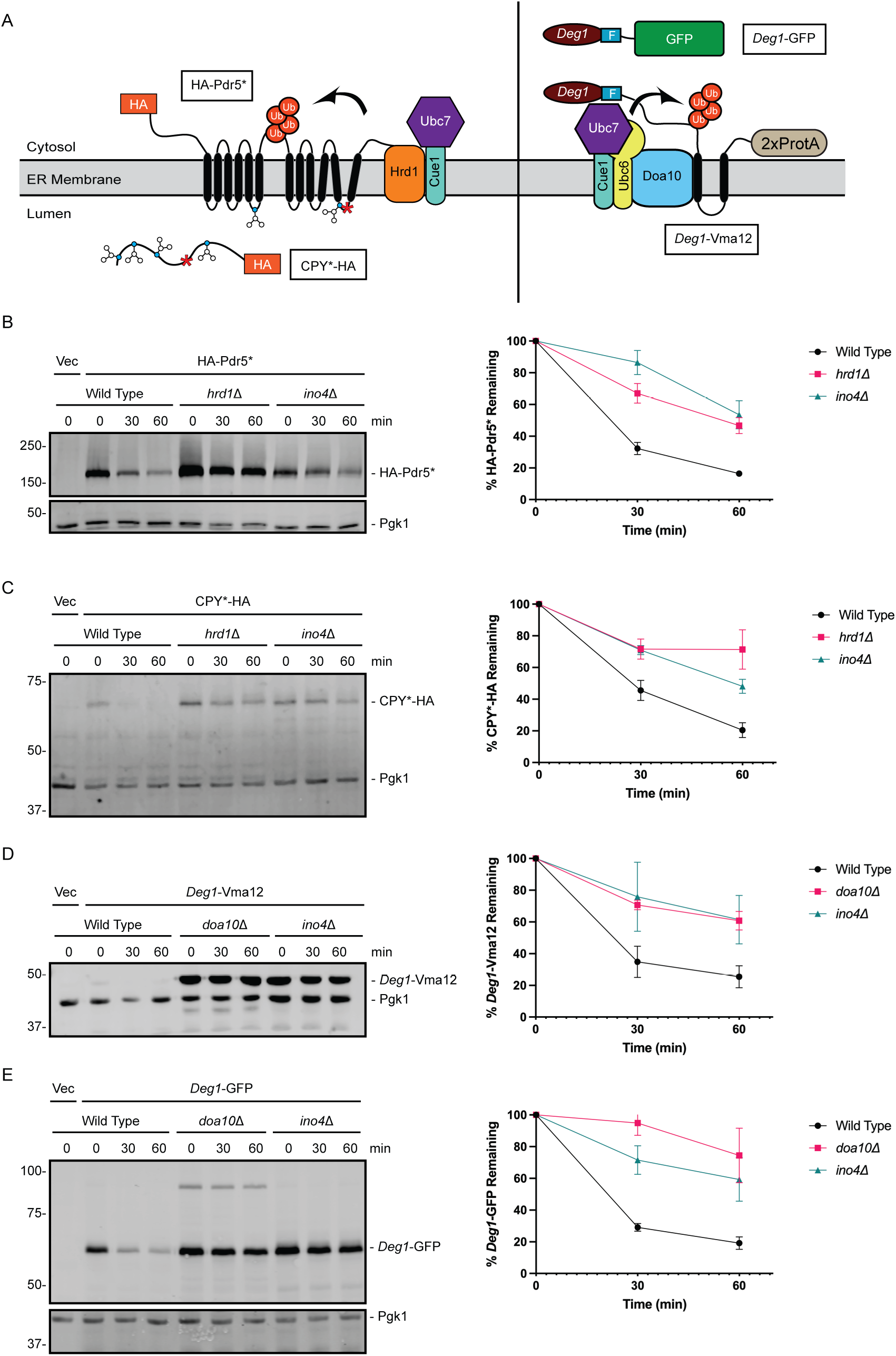
*INO4* deletion impairs ERAD of Hrd1 and Doa10 substrates. **(A)** ERAD substrates of Hrd1 and Doa10 analyzed in this figure. F, Flag; ProtA, Protein A; Ub, ubiquitin **(B - E)** Left, Yeast of the indicated genotypes were transformed with a plasmid encoding indicated ERAD substrates or an empty vector and subjected to cycloheximide chase and western blot analysis to detect the ERAD substrate and Pgk1. Right, quantification of experiments. Error bars represent Standard Error of the Mean of n = 3-5 experiments.

We analyzed abundance of plasmid-encoded Hrd1-3HA in yeast expressing or lacking *INO4*. Hrd1-3HA is functional *in vivo* (Sato *et al*. 2009). Hrd1 abundance was elevated in *ino4*Δ yeast, but not significantly (**Figure 5A**). Both Doa10 and Hrd1 require the ubiquitin-conjugating enzyme Ubc7 and its membrane anchor Cue1. Under some circumstances, Hrd1 undergoes autoubiquitylation and degradation in a Ubc7-dependent manner (Plemper *et al*. 1999; Gardner *et al*. 2000). Thus, impaired ERAD and increased Hrd1 abundance could be attributed to reduced abundance of Ubc7. We therefore assessed abundance of 2HA-tagged Ubc7 in wild type yeast, yeast lacking Cue1, and yeast lacking Ino4. HA-tagging of Ubc7 does not abolish activity or ubiquitin ligase interaction ((Ravid and Hochstrasser 2007; Kreft and Hochstrasser 2011). Consistent with previous reports demonstrating *CUE1* deletion destabilizes Ubc7 (Biederer *et al*. 1997; Gardner *et al*. 2001; Ravid and Hochstrasser 2007), steady state abundance of Ubc7-2HA was significantly reduced in *cue1*Δ cells (**Figure 5B**). Ubc7-2HA abundance was similarly reduced in *ino4*Δ yeast. Thus, one potential mechanism by which *INO4* deletion compromises ERAD is through reduction of abundance of one or more components of the ubiquitylation machinery, such as Ubc7.

**Figure 5.**
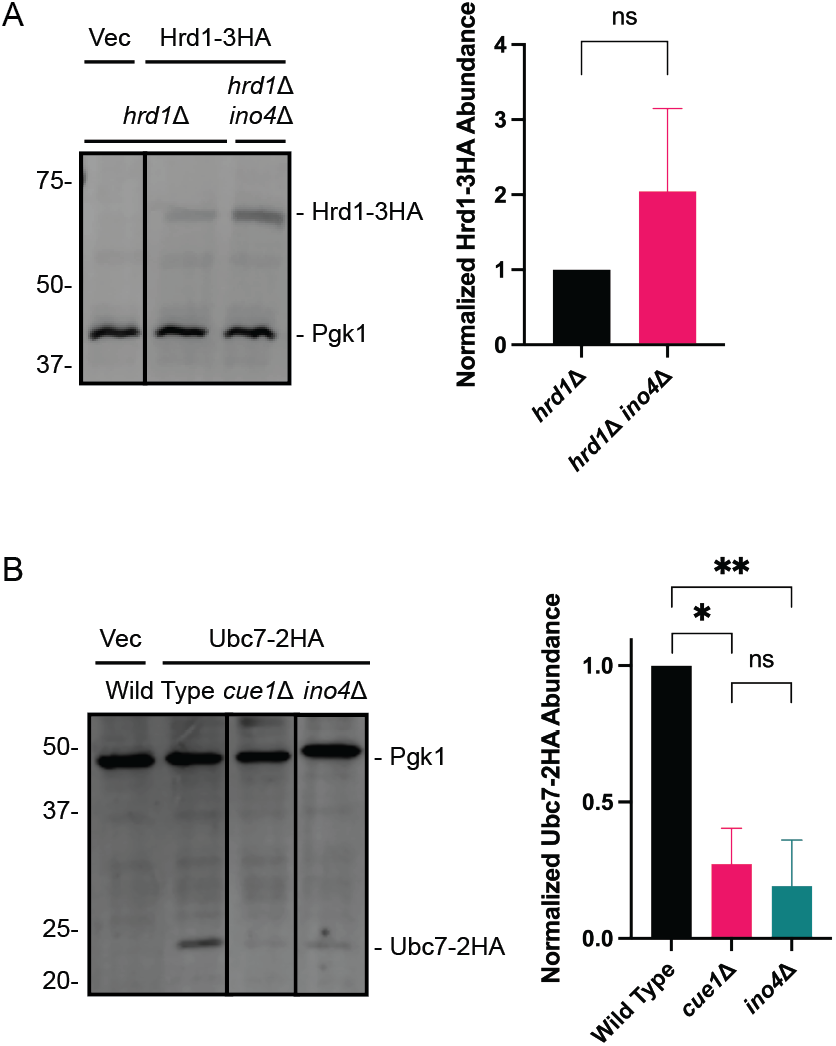
Loss of *INO4* causes a reduction in Ubc7-2HA abundance. **(A and B)** Left, Yeast of the indicated genotypes were transformed with a plasmid encoding Hrd1-3HA, Ubc7-2HA, or an empty vector, harvested, lysed, and subjected to anti-HA and anti-Pgk1 western blotting. Right, quantification of experiments. Error bars represent Standard Error of the Mean of n = 3 experiments. Means in **(A)** were evaluated by a two-tailed *t*-test. Means in **(B)** were evaluated by one-way ANOVA with a Tukey *post hoc* test. *, p < 0.001; **, p < 0.05. ns, not significant.

### Genetic perturbation of lipid biosynthesis sensitizes yeast to hygromycin B

Given the broad requirement of *INO4* for degradation of several aberrant proteins at the ER membrane, we predicted disruption of *INO4* and other genes required for lipid biosynthesis would sensitize yeast to conditions associated with an elevated burden of aberrant or misfolded proteins. We analyzed growth of wild type yeast and yeast possessing deletions or hypomorphic alleles of *HRD1, INO4, CDS1, CHO1* (required for phosphatidylserine, phosphatidylethanolamine, and phosphatidylcholine synthesis), *INO1*, or *ERG1* in the presence of hygromycin B (**Figure 6A**), tunicamycin (**Figure 6B**), or a range of temperatures (**Figure 6C**). Hygromycin B distorts the ribosome aminoacyl site, resulting in globally increased production of aberrant polypeptides (Brodersen *et al*. 2000; Ganoza and Kiel 2001). Tunicamycin inhibits N-linked glycosylation, specifically impairing ER proteostasis (Travers *et al*. 2000).

**Figure 6.**
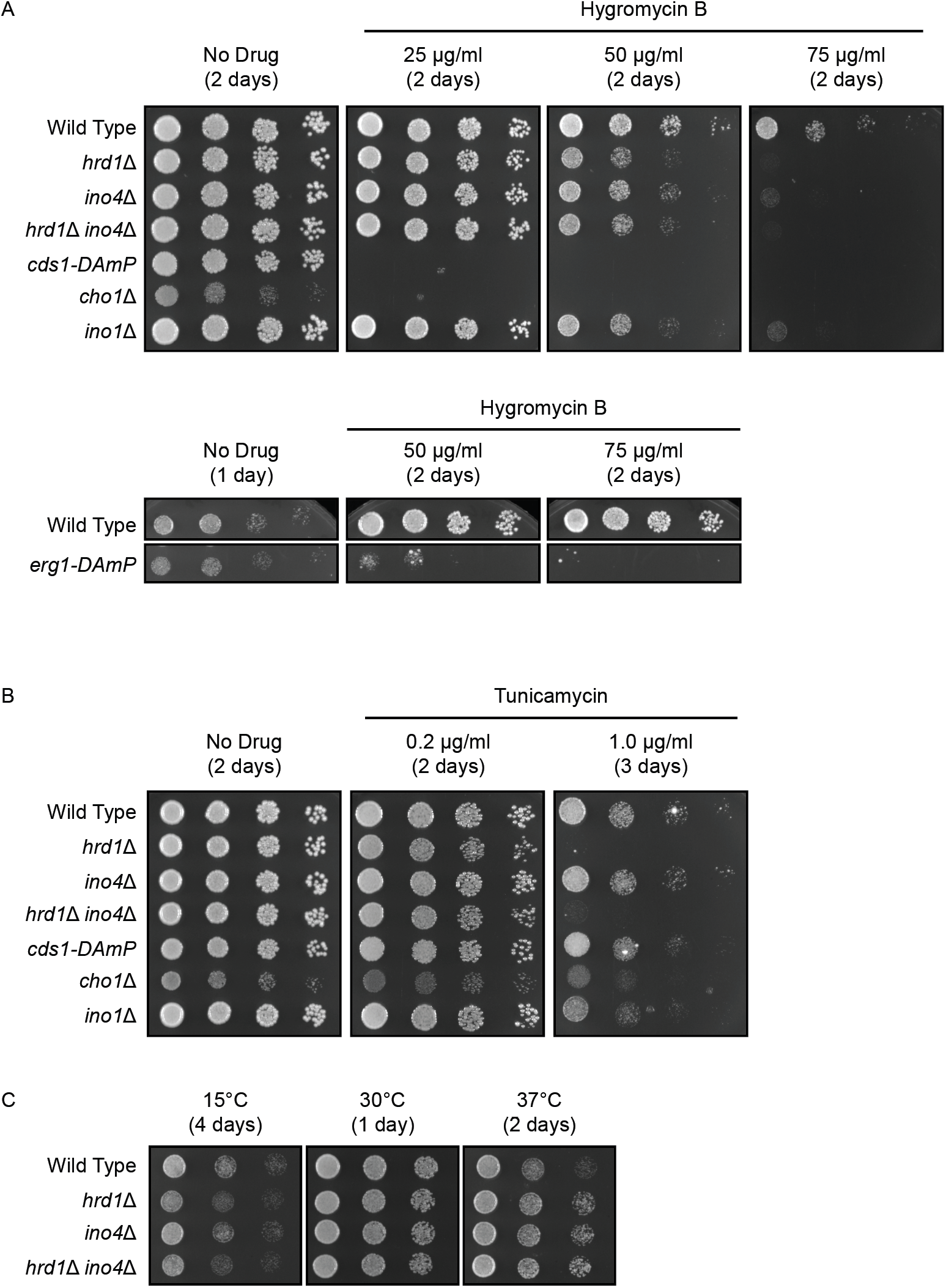
Stress sensitivity of yeast defective for lipid biosynthesis. Yeast of the indicated genotypes were serially diluted and spotted onto agar plates containing the indicated medium. Plates were incubated at 30°C or the indicated temperatures and imaged after 1-3 d.

As previously documented, *HRD1* deletion strongly sensitized yeast to hygromycin B and tunicamycin (Kapitzky *et al*. 2010; Crowder *et al*. 2015; Niekamp *et al*. 2019; Runnebohm *et al*. 2020a; Woodruff *et al*. 2021). *INO4* deletion and mutation of genes regulated by *INO4* sensitized yeast to hygromycin B, but not tunicamycin. *hrd1*Δ *ino4*Δ yeast did not exhibit enhanced sensitivity to either compound relative to *hrd1*Δ yeast. Yeast lacking *HRD1* and *INO4* (or both) did not exhibit sensitivity to reduced (15°C) or elevated (37°C) temperatures. Sensitivity of *ino4*Δ, *cds1-DAmP, cho1*Δ, *ino1*Δ, and *erg1-DAmP* yeast to hygromycin B is consistent with a requirement of membrane lipid homeostasis for proteostasis.

## DISCUSSION

In this study, we conducted a screen to identify novel genetic requirements for degradation of a model translocon-associated Hrd1 ERAD substrate. This screen revealed three genes (*INO4, KAR3*, and *SET2*) required for efficient *Deg1*\*-Sec62 degradation. We characterized the involvement of *INO4* and lipid biosynthetic enzymes in ERAD. *KAR3* was previously identified in a screen for genes required for degradation of a Doa10 substrate (Ravid *et al*. 2006). Further, gene ontology analysis yielded an enrichment of genes linked to sulfur metabolism. The impact of *KAR3* deletion on ERAD and the interplay between sulfur metabolism and ER homeostasis will be explored in subsequent studies.

To our knowledge, neither *INO2* nor *INO4* have been identified in previous screens for genetic requirements of ER protein degradation (e.g. (Hampton *et al*. 1996; Knop *et al*. 1996; Swanson *et al*. 2001; Ravid *et al*. 2006; Kohlmann *et al*. 2008; Ast *et al*. 2016; Neal *et al*. 2018)). This is likely related to the fact that yeast lacking *INO2* or *INO4* possess a decreased growth rate in minimal media (**Figure S2**), commonly used in growth reporter-based genetic screens. Our identification of *INO4* highlights the power of our screen to reveal genetic requirements for degradation.

Our results indicate ERAD is broadly sensitive to perturbations in lipid homeostasis. Deletion of either gene encoding members of the Ino2/Ino4 transcriptional regulator stabilizes *Deg1*\*-Sec62 to a similar extent as *HRD1* deletion. Stabilization is likely due to altered membrane lipid composition, as supplementation with lipid biosynthetic intermediates rescued the *Deg1*\*-Sec62 degradation defect of *ino4*Δ yeast. Further, disruption of Ino2/Ino4-regulated genes encoding lipid biosynthetic enzymes impede degradation, as does perturbation of a gene encoding an enzyme required for sterol biosynthesis (*ERG1*, not regulated by Ino2/Ino4).

This work builds on an expanding literature linking lipid homeostasis to ER proteostasis. Consistent with broad conservation of the relationship between lipid and protein homeostasis, inhibition of long-chain acyl-coA synthetases impairs glycan trimming, ER extraction, and degradation of a subset of glycosylated substrates of the mammalian Hrd1 ubiquitin ligase (To *et al*. 2017). Additionally, degradation of the misfolded ER luminal yeast protein CPY* is sensitive to *CHO2* deletion (Thibault *et al*. 2012) and, more modestly, to mutations that impair lipid droplet formation (Yap *et al*. 2020). Yeast and mammalian homologs of Hrd1 and Doa10 promote feedback-regulated degradation of sterol-biosynthetic enzymes as well as turnover of proteins implicated in triacylglycerol and low-density-lipoprotein synthesis (Hampton *et al*. 1996; Garza *et al*. 2009; Jo *et al*. 2011; Foresti *et al*. 2013; Stevenson *et al*. 2016; Huang and Chen 2023). Moreover, the ER-resident rhomboid pseudoprotease Dfm1 was recently shown to regulate the ER export and Golgi-associated degradation of Orm2, an inhibitor of sphingolipid biosynthesis (Bhaduri *et al*. 2022). Lipid bilayer stress activates the yeast and mammalian ER unfolded protein response (UPR) via a distinct mechanism than activation by unfolded proteins (Jonikas *et al*. 2009; Promlek *et al*. 2011; Thibault *et al*. 2012; Volmer *et al*. 2013; Halbleib *et al*. 2017; Ho *et al*. 2020; Xu and Taubert 2021). The protein homeostatic machinery induced by the UPR buffers the toxic effects of disrupted lipid homeostasis (Thibault *et al*. 2012), and genes required for lipid biosynthesis are synthetically lethal with those encoding UPR mediators (Costanzo *et al*. 2010; Thibault *et al*. 2012).

Our results extend the impact of altered membrane composition to include several different classes of ERAD substrates of two different ubiquitin ligases. *INO4* deletion stabilized a panel of soluble and transmembrane substrates of two ERAD ubiquitin ligases, Hrd1 and Doa10, and a translocon-associated substrate of Hrd1. Further, deletion of *INO4* and mutation of genes required for different aspects of lipid biosynthesis sensitized yeast to hygromycin B, which is expected to increase the cellular burden of misfolded proteins. Future studies will be conducted to determine whether perturbed lipid biosynthesis impairs degradation by ubiquitin ligases in other cellular compartments (e.g. inner nuclear membrane Asi complex).

How does disrupted lipid composition impair ER protein quality control? As a consequence of altered membrane fluidity or protein-lipid interactions, perturbation of membrane lipid composition may change the abundance, structure, membrane integration or docking, or localization of substrate recognition or ubiquitylation machinery. Our results indicate the ubiquitin-conjugating enzyme Ubc7 (which is required for both Hrd1-and Doa10-dependent ERAD) is present in reduced abundance in *ino4*Δ yeast. Ubc7 is anchored to the ER membrane – and stabilized – by Cue1 (Biederer *et al*. 1997; Gardner *et al*. 2001; Ravid and Hochstrasser 2007). Recent work has demonstrated that altered membrane lipid composition modestly destabilizes Cue1 (Shyu *et al*. 2019). Reduced Ubc7 expression (via either accelerated degradation or dampened rate of synthesis) might contribute to broad ERAD impairment.

*Deg1*\*-Sec62 is glycosylated upon its aberrant translocon engagement (Rubenstein *et al*. 2012). This modification is delayed in yeast lacking several genes encoding lipid biosynthetic enzymes, consistent with dampened translocation rate. While it remains possible that lipid bilayer stress slows ER import of one or more proteins required for ERAD, CPY*, which undergoes glycosylation upon ER import, does not exhibit altered mobility in *ino4*Δ yeast. This suggests *INO4* deletion does not cause a generalized translocation block. It is also conceivable that altered membrane lipid composition impedes ERAD substrate retrotranslocation (movement from the ER to the cytosol). However, impaired retrotranslocation cannot explain the totality of the impact of *INO4* deletion, as a model soluble, cytosolic Doa10 substrate (*Deg1*-GFP) was stabilized in *ino4*Δ yeast.

In contrast to impaired degradation of the Doa10 substrates evaluated in this study, lipid bilayer stress caused by phosphatidylcholine depletion (i.e. *OPI3* deletion) accelerates degradation of Doa10 substrate Sbh1 (Shyu *et al*. 2019). Sbh1 is atypically degraded by Doa10, as its turnover occurs independently of cytosolic lysine residues (Shyu *et al*. 2019). This suggests that Doa10 activity *per se* may not be impaired by phosphatidylcholine deficiency, as is caused by *INO4* deletion (Loewy and Henry 1984). Comparing the mechanism of canonical versus atypical Doa10 degradation mechanisms may reveal molecular factors that are differentially sensitive to membrane lipid composition.

In a previous study, we found *Deg1*\*-Sec62 degradation occurs with wild type kinetics in the context of inositol depletion (Buchanan *et al*. 2019). Data from the present study appear to contract this observation, which we reproduced (**Figure S3**). We speculate this disparity reflects differences in duration of lipid perturbation. In our earlier study, yeast experienced inositol restriction for five h, whereas yeast in the present study were subjected to genetic (i.e. long-term) perturbations in lipid biosynthesis.

Perturbed membrane lipid composition has been implicated in multiple diseases, including non-alcoholic fatty liver disease (Li *et al*. 2006; Arendt *et al*. 2013), obesity and type II diabetes (Kim *et al*. 2016), muscular dystrophy (Mitsuhashi and Nishino 2011), and cardiomyopathies (Bargui *et al*. 2021). Our results suggest alterations in lipid profiles associated with these disease states are likely to impair ERAD. Phospholipid metabolism may also be altered in Alzheimer’s disease, (Frisardi *et al*. 2011; Grimm *et al*. 2011; Kosicek and Hecimovic 2013; Tajima *et al*. 2013; Emre *et al*. 2021). The involvement of ERAD in Alzheimer’s disease progression is complex; multiple physiological ERAD substrates differentially impact amyloid-Δ accumulation (Tanabe *et al*. 2012; Zhu *et al*. 2017). Further, deletion of several genes linked to lipid metabolism or transport have been found to alter intracellular localization, aggregation, and toxicity of amyloid-Δ in a yeast cellular model for Alzheimer’s disease (Nair *et al*. 2014; Chen *et al*. 2020). Further investigation of the relationship between lipid and protein homeostasis may inform improved understanding and treatment of diseases associated with disruptions in cellular lipid dynamics.

## ACKNOWLEDGMENTS

We thank Jacob Davis, Molly Dolan, Jack Elo, Seth Horowitz, Brianne McCord, Ashleigh South, Lauren Huffman Wade, and Sheldon Watts for lab assistance and assistance with generating reagents used for this study. Pilot experiments to determine the abundance of *Deg1*\*-Sec62 in mutants identified in our screen were performed by undergraduate students in the Fall 2017, Spring 2018, Fall 2018, Spring 2019, Fall 2019, and Spring 2020 sections of Methods in Cell Biology (BIO 315) course in the Ball State University Department of Biology. We thank Teaching Assistants Melissa Evans, Jacob Davis, Adam Richardson, and Abigail Scott for assisting BIO 315 students in these pilot experiments. We thank Ashley Kalinski, Douglas Bernstein, Jason True, Susan McDowell, Heather Bruns, Jennifer Metzler, Mark Hochstrasser, Stefan Kreft, Tommer Ravid, Oliver Kerscher, and Adrian Mehrtash for stimulating conversations and suggestions for experiments. We thank Stefan Kreft (University of Kontsanz), Nydia Van Dyk (University of Toronto), Charlie Boone (University of Toronto), Randy Hampton (University of California San Diego), James Olzmann (University of California Berkeley), Dieter Wolf (University of Stuttgart), and Mark Hochstrasser (Yale University) for plasmids, strains, and antibodies. We remain grateful to *Saccharomyces Genome Database* (SGD; yeastgenome.org) for curating yeast genetic information; without SGD, this work would be immeasurably more difficult.

## FIGURES AND FIGURE LEGENDS

**Figure S1.**
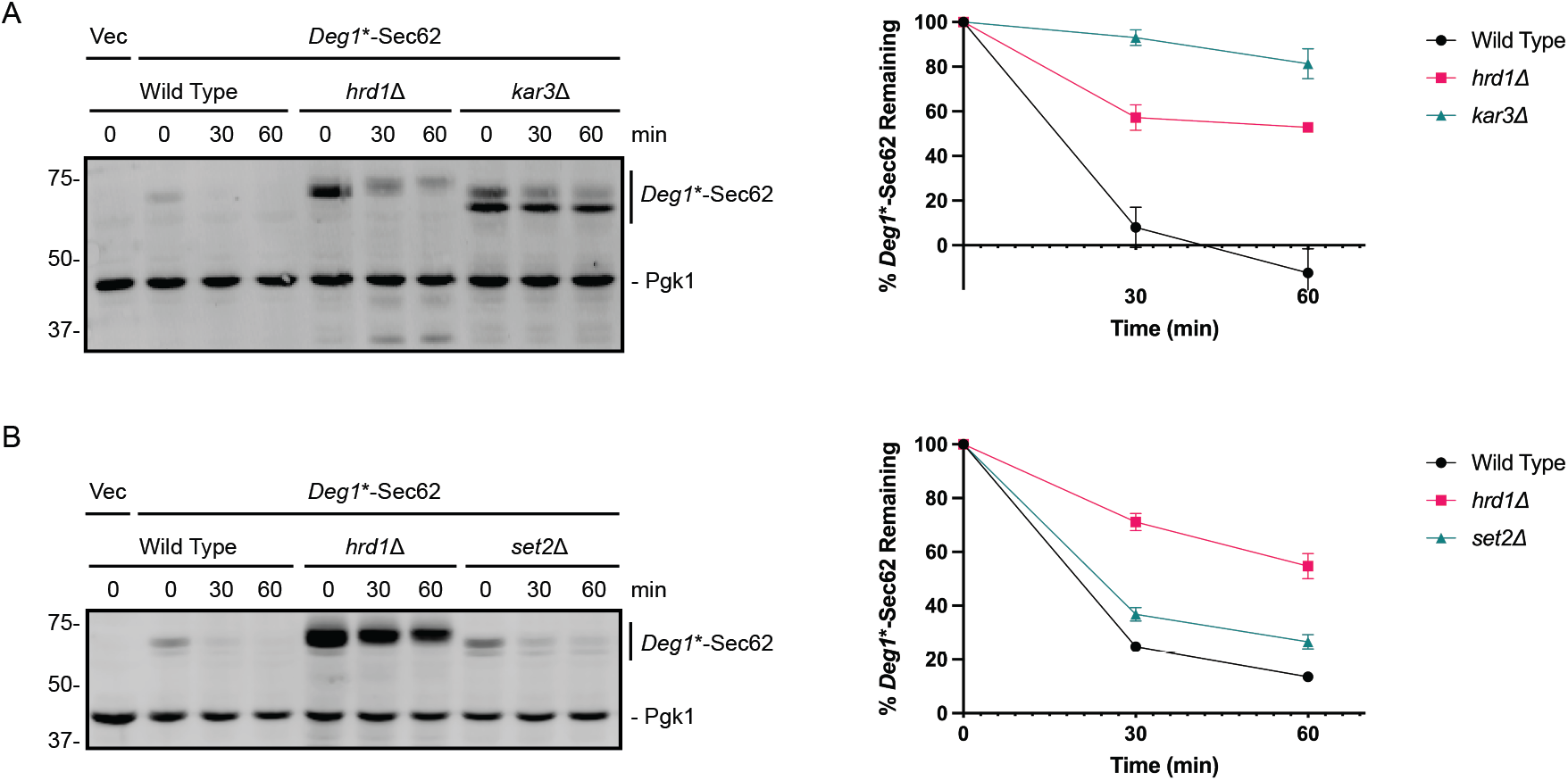
Stabilization of *Deg1*\*-Sec62 in yeast lacking *KAR3* and *SET2*. Left, Yeast of the indicated genotypes were transformed with a plasmid encoding *Deg1*\*-Sec62 or an empty vector and subjected to cycloheximide chase and western blot analysis to detect *Deg1*\*-Sec62 and Pgk1. Right, quantification of experiments. Error bars represent Standard Error of the Mean of n = 3 experiments.

**Figure S2.**
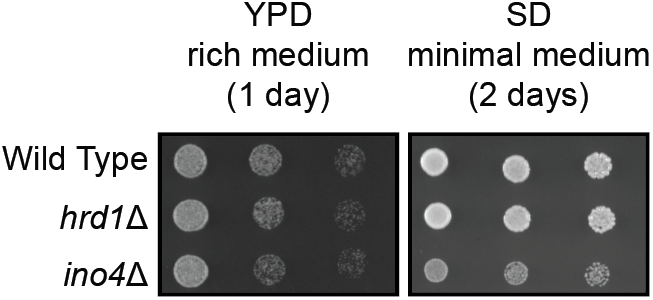
Yeast lacking *INO4* exhibit growth defect on minimal medium. Yeast of the indicated genotypes were serially diluted and spotted onto agar plates containing YPD (Yeast Extract-Peptone-Dextrose rich medium) or SD (Synthetic Defined minimal medium). Plates were incubated at the indicated temperatures and imaged after 1-2 d.

**Figure S3.**
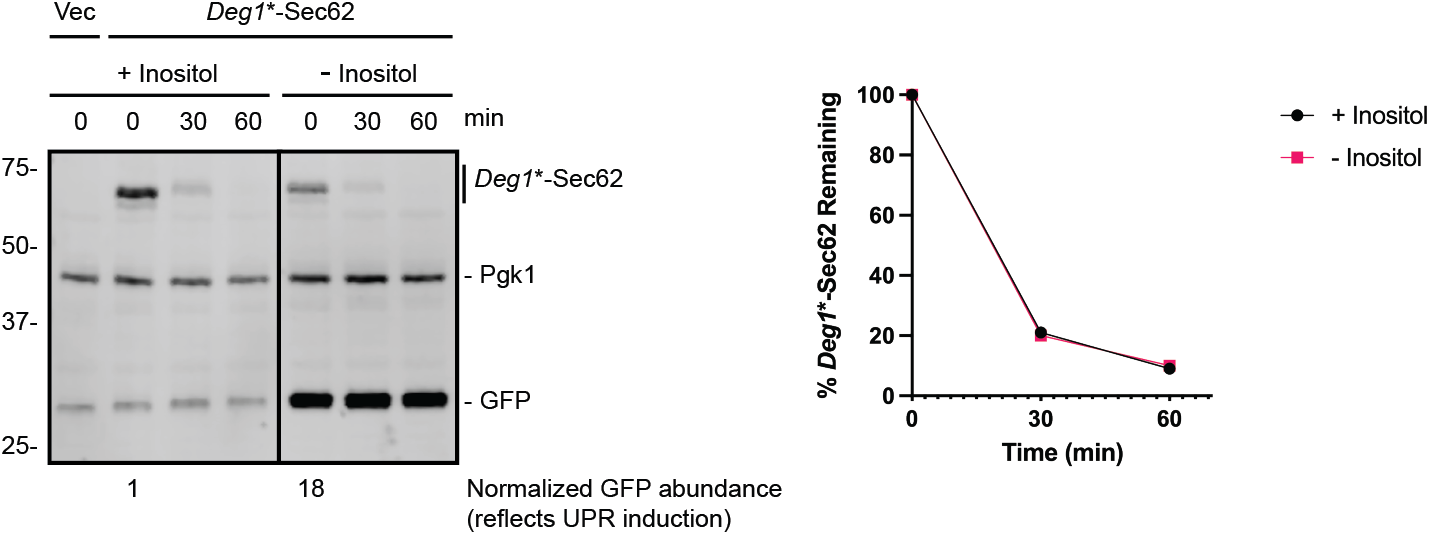
Short-term inositol limitation does not impair *Deg1*\*-Sec62 degradation. Wild type yeast were transformed with a plasmid encoding *Deg1*\*-Sec62 or an empty vector, cultured to mid-exponential phase, washed six times in medium with inositol (+ inositol) or lacking inositol (-inositol), resuspended in the same media, and subjected to cycloheximide chase and western blot analysis to detect *Deg1*\*-Sec62 and Pgk1. Right, quantification of experiment.

## SUPPLEMENTARY TABLES

**Table S1.**
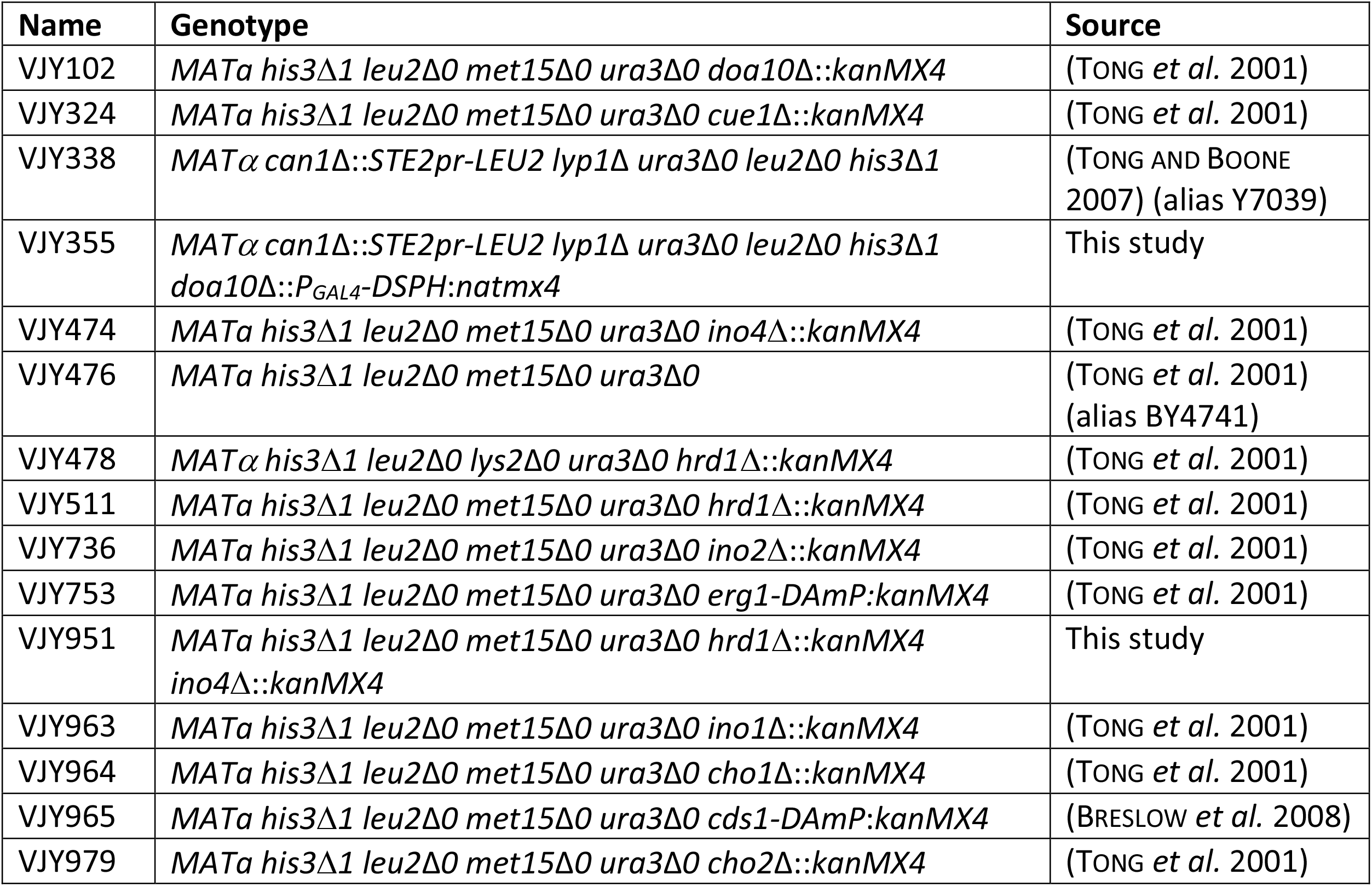
Yeast strains used in this study.

**Table S2.** Plasmids used in this study.

| Name | Alias | Yeast selection marker | Yeast plasmid type | Description | Source |
| --- | --- | --- | --- | --- | --- |
| pHA-Pdr5* | pVJ1;<br>pRH2312 | *HIS3* | CEN | HA-tagged Pdr5*; Pdr5* = C1427Y | (SATO *et al.* 2009) |
| YCp50-*P<sub>PRC1</sub>*-CPY*-HA | pVJ2;<br>pDN431 | *URA3* | CEN | HA-tagged CPY* driven by native promoter; CPY* = G255R | (NG *et al.* 2000) |
| pRS313 | pVJ26 | *HIS3* | CEN | Empty vector | (SIKORSKI AND HIETER 1989) |
| pRS316 | pVJ27 | *URA3* | CEN | Empty vector | (SIKORSKI AND HIETER 1989) |
| P416-*Deg1*-GFP2 | pVJ205;<br>pUL28 | *URA3* | CEN | *Deg1* fused to two copies of GFP | (LENK AND SOMMER 2000) |
| p416- *P<sub>MET25</sub>* - *Deg1* *-Sec62-2xProtA | pVJ317 | *URA3* | CEN | *Deg1* *-Flag-Sec62-2xProtA driven by *MET25* promoter; *Deg1* * = F18S, I22T | (RUBENSTEIN *et al.* 2012) |
| p416-*P<sub>GPD</sub>*-*Deg1*-Vma12-2xProtA | pVJ343 | *URA3* | CEN | *Deg1*-Flag-Vma12-2xProtA ("*Deg1*-Vma12") driven by *TDH3* (*GPD*) promoter | (BUCHANAN *et al.* 2019) |
| p414-*P<sub>GAL4</sub>*-*Deg1*-Sec62-2xProtA-His3:natmx4 | pVJ490 | *TRP1*,<br>*natMX4* | CEN | Source of *P<sub>GAL4</sub>*-*Deg1*-Sec62-His3:natmX4 cassette to generate query strain for screen;<br>*Deg1*-Flag-Sec62-2xProtA-His3 ("*Deg1*-Sec62-His3") driven by *GAL4* promoter; *natMX4* is a distinct gene with its own promoter and terminator | This study |
| pRS313-*ADH*-Ubc7-2HA | pVJ520;<br>STK06-2-5 | *HIS3* | CEN | 2HA-tagged Ubc7 driven by *ADH* promoter | (STÜRNER 2015) |
| pRS314-UPRE-GFP | pVJ552 | *TRP1* | CEN | GFP driven by Unfolded Protein Response Element (UPRE) | (OLZMANN AND KOPITO 2011) |

**Table S3.**
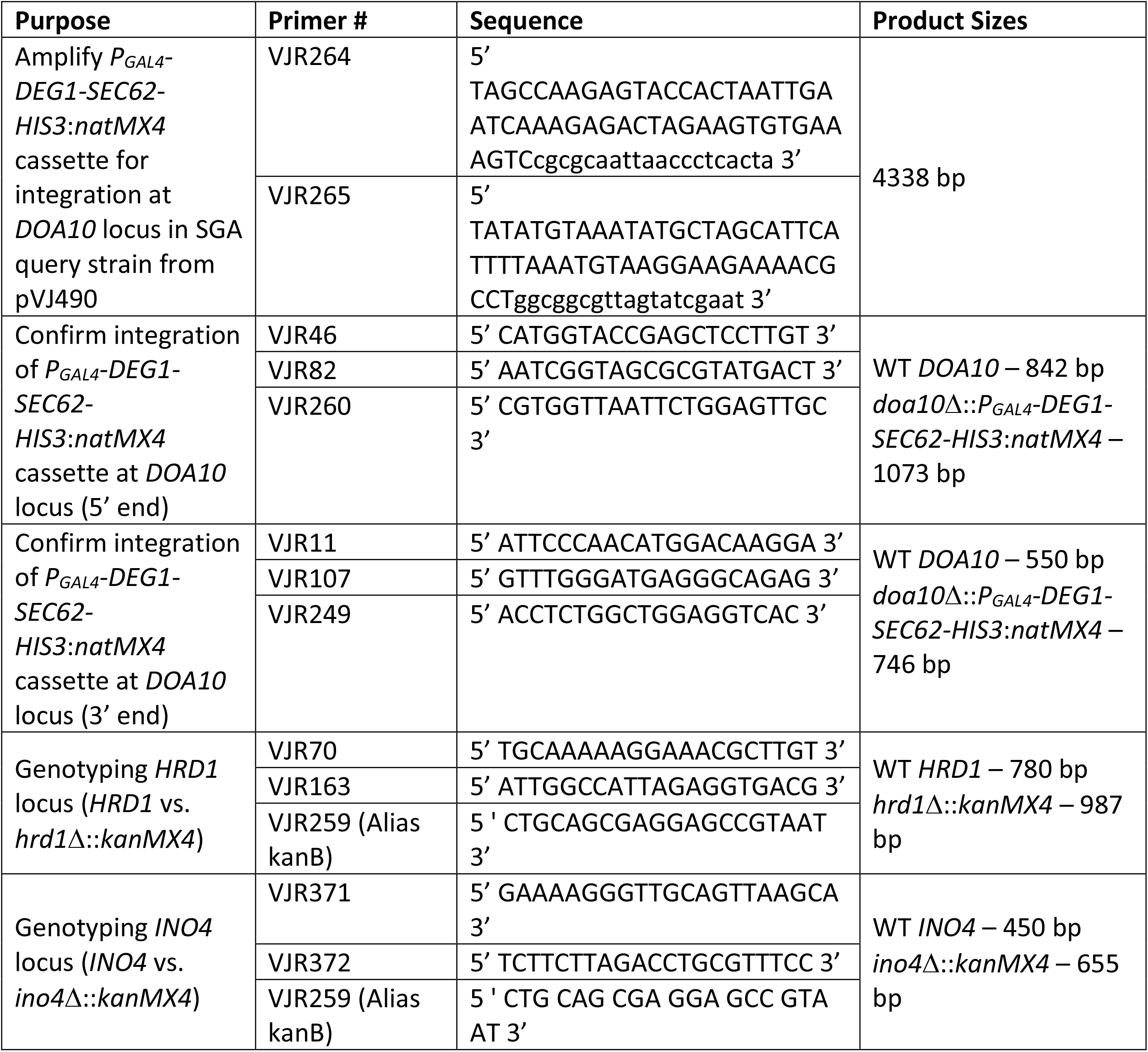
Primers used in this study.

**Table S4.** Outline of SGA Screen Procedure.

| Description | Media | Agar or liquid media | Time |
| --- | --- | --- | --- |
| Prepare a lawn of query strain (VJY355) | YPD | Agar | 2 d |
| Pin array of library strains from 96-well plate of cryopreserved yeast | YPD | Agar | 2 d |
| Mate yeast by replica-pinning query strain and library strains onto fresh plate | YPD | Agar | 1 d |
| Mated diploid selection | YPD + 100 µg/ml nourseothricin (clonNAT; WERNER BioAgents) + 200 µg/ml G418 (G418 Sulfate; CALBIOCHEM) | Agar | 2 d |
| Sporulation | 1% Potassium Acetate, 0.1% yeast extract, 0.05% glucose, 0.002% adenine, 0.004% uracil, 0.0005% arginine, 0.00025% histidine, 0.0015% isoleucine, 0.0015% leucine, 0.001% lysine, 0.00025% methionine, 0.0015% phenylalanine, 0.00125% threonine, 0.001% tryptophan | Agar | 5-7 d |
| MAT<sub>a</sub> selection | 0.67% yeast extract without amino acids, 2% glucose, 0.0005% arginine, 0.00025% histidine, 0.0015% isoleucine, 0.0015% leucine, 0.001% lysine, 0.00025% methionine, 0.0015% phenylalanine, 0.00125% threonine, 0.001% tryptophan, 0.004% adenine, 0.008% uracil, 50 µg/ml canavanine; 50 µg/ml thialysine | Agar | 2 d |
| Gene deletion or hypomorphic allele selection | MAT<sub>a</sub> selection media + 200 µg/ml G418 | Agar | 2 d |
| Selection of deletion or hypomorphic allele *AND* *Deg1-Sec62-His3:natMX4* reporter | MAT<sub>a</sub> selection media + 200 µg/ml G418 + 100 µg/ml nourseothricin | Agar | 2 d |
| Prepare 96-well plates with liquid cultures | MAT<sub>a</sub> selection media + 200 µg/ml G418 + 100 µg/ml nourseothricin | Liquid | 2 d |
| Transfer 40 µl of liquid culture to fresh media to | MAT<sub>a</sub> selection media + 200 µg/ml G418 + 100 µg/ml nourseothricin *without* histidine | Liquid | 11 h<br>(Record |
| select for genes required<br>for *Deg1*-Sec62-His3<br>degradation |  |  | OD<sub>595</sub> at<br>beginning<br>and end of<br>11 h) |

All steps performed at 30°C, except for mating and sporulation, which were performed at 23°C.
All transfers (except final transfer to screen media) performed using sterile 96-pronged pinners.

**Table S5.** Gene Ontology Process Term analysis for genes identified in genome-wide screen.

| **Gene Ontology term** | **Cluster frequency** | **Genome frequency** | **Corrected p value** | **Genes** |
| --- | --- | --- | --- | --- |
| sulfate assimilation | 5 of 128 genes, 3.9% | 9 of 7166 genes, 0.1% | 0.00012 | *MET3 MET5 MET8 MET10 MET14* |
| sulfur compound biosynthetic process | 10 of 128 genes, 7.8% | 82 of 7166 genes, 1.1% | 0.00102 | *HOM2 HOM3 MET2 MET3 MET5 MET6 MET10 MET14 PDA1 SAM1* |
| sulfur amino acid metabolic process | 8 of 128 genes, 6.2% | 49 of 7166 genes, 0.7% | 0.00124 | *HOM2 HOM3 MET2 MET3 MET5 MET6 MET14 SAM1* |
| hydrogen sulfide metabolic process | 4 of 128 genes, 3.1% | 8 of 7166 genes, 0.1% | 0.00395 | *MET3 MET5 MET10 MET14* |
| hydrogen sulfide biosynthetic process | 4 of 128 genes, 3.1% | 8 of 7166 genes, 0.1% | 0.00395 | *MET3 MET5 MET10 MET14* |

128 genes with $\Delta OD_{595}$ values greater than or equal to 0.079 in screen for genes with potential roles in ER protein degradation were analyzed using the Gene Ontology Term Finder at the Saccharomyces Genome Database (https://www.yeastgenome.org/goTermFinder) using a p-value cutoff of 0.01. No significant Function or Component GO Terms were found the input list of 128 genes.

## SUPPLEMENTAL FILES

**Supplemental File 1. Results of genome-wide screen**.

